## Supplementary Data for "Adenovirus maturation establishes the transcription competent packaging of its genome"

* corresponding authors

| Genotype | Name | Treatment | number of reads | alignment rate | aligned reads | coverage |
|---|---|---|---|---|---|---|
| WT | free DNA R1 | DMS | 2.70E+07 | 3.28% | 8.22E+05 | 9.89E+02 |
| WT | free DNA R2 | DMS | 3.20E+07 | 2.59% | 7.76E+05 | 9.35E+02 |
| WT | Ad-wt R1 | DMS | 3.30E+07 | 99.71% | 3.04E+07 | 3.66E+04 |
| WT | Ad-wt R2 | DMS | 2.40E+07 | 99.42% | 2.23E+07 | 2.68E+04 |
| ts1 | Ad-*ts1* R1 | DMS | 2.60E+07 | 93.39% | 2.21E+07 | 2.66E+04 |
| ts1 | Ad-*ts1* R2 | DMS | 2.90E+07 | 93.50% | 2.55E+07 | 3.06E+04 |

**Table 1: Overview to the samples, read depth and annotation rate.**

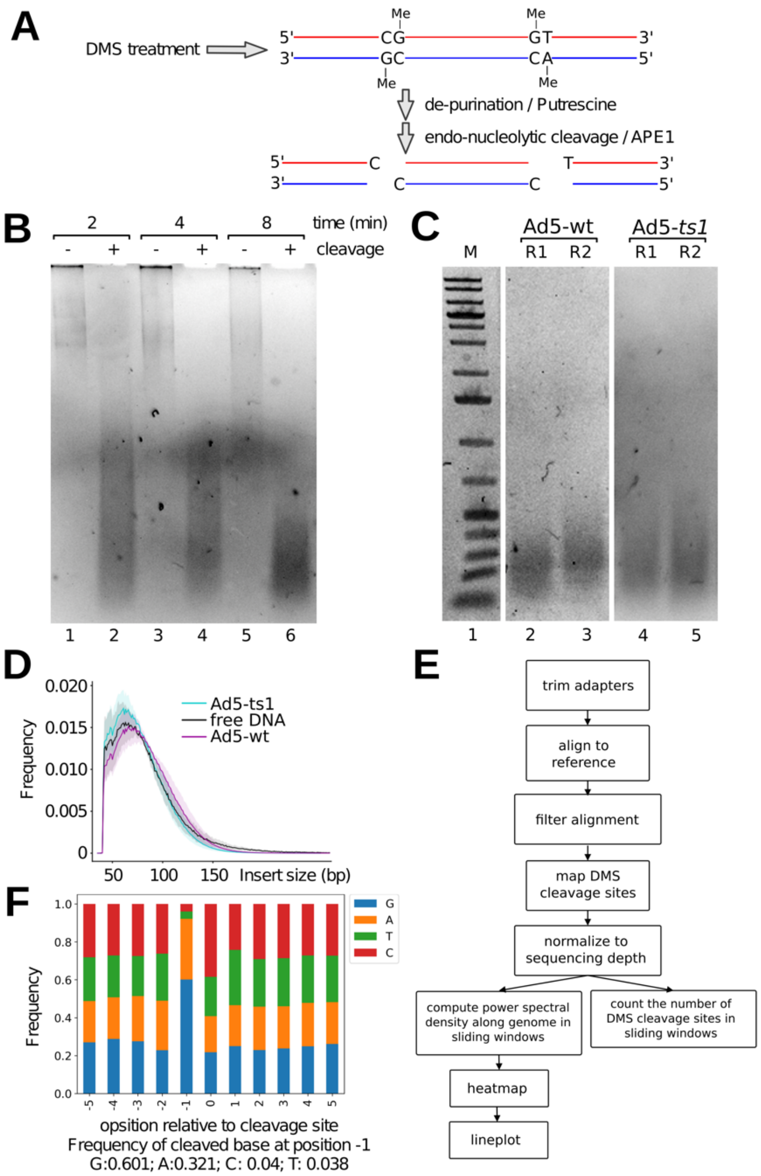


**Figure S1: Depiction of DMS-seq workflow and controls.** (A) Scheme showing the DMS caused fragmentation of DNA by Putrescine and APE1 (B) Agarose gel (1.3%) of viral DNA treated in the absence or presence of 5% DMS for the indicated time points. Viruses were treated with DMS as shown, and the purified DNA was cleaved in the presence of Putrescine and Ape1. (C) Viral DNA fragmented after treatment with 2% DMS for 4 minutes and subsequent cleavage reaction was used for library preparation (D) Fragment size distribution histogram of free DNA, Ad-wt and Ad-ts1 (E) Overview to the pipeline for the processing of DMS-seq data (F) Plot of nucleotide frequencies relative to fragment cleavage positions, as indicated.

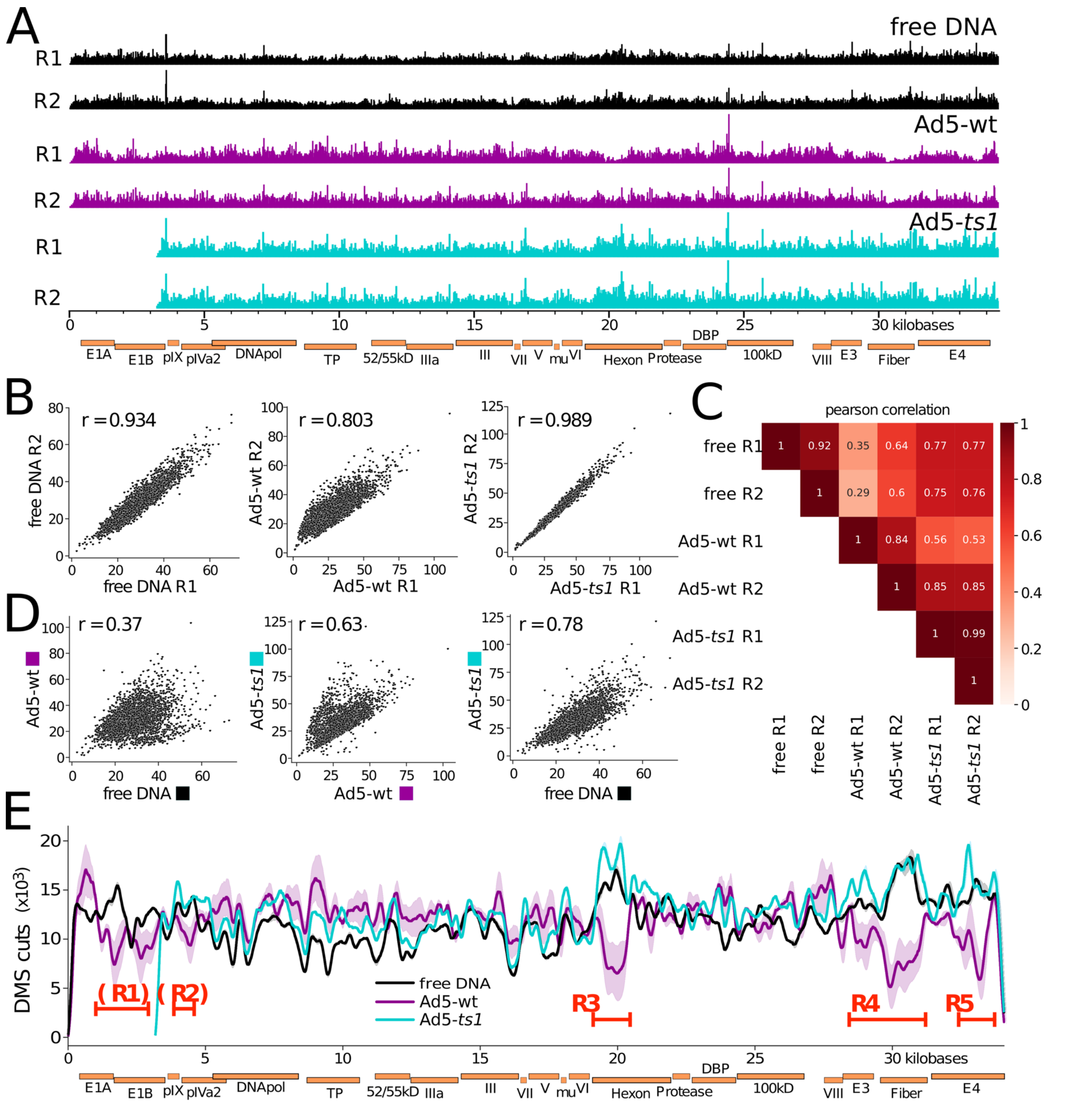


**Figure S2: DMS cleavage site distribution along genome** (A) Bar chart of cleavage sites along the reference genomes, individually displayed for each replicate (B) Correlation between replicates in 10 bp bins along the genome (C) Correlation Matrix for the individual samples generated by deeptools (bin size 5 bp; see Materials and Methods) (D) Correlation between samples (replicates combined) in 10 bp bins along the genome (E) Count of DMS cut sites in a 400 bp sliding window with 100 bp step size across the reference genomes.

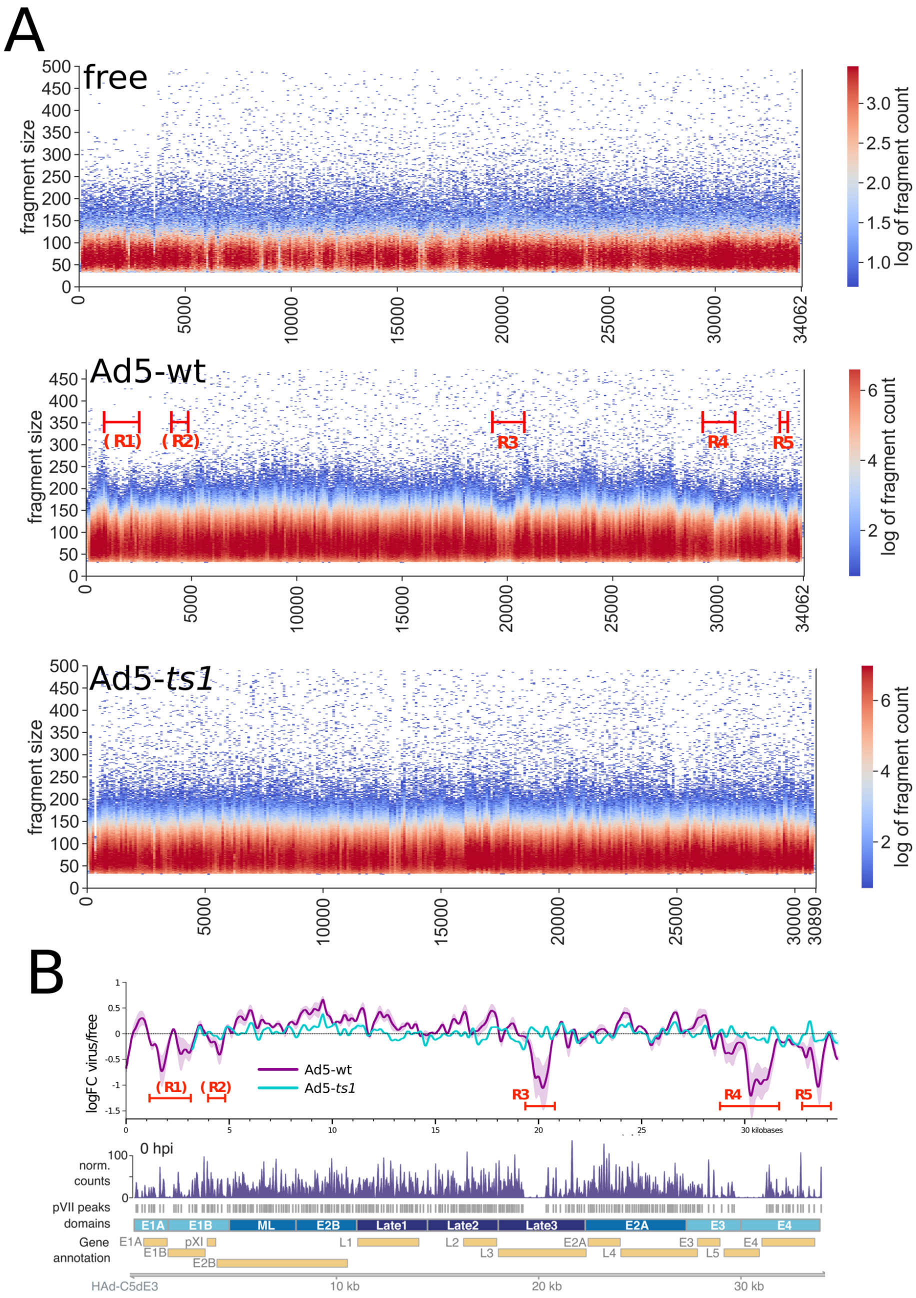


**Figure S3:** (A) V-plots, plotting the fragment size against the fragment mid-point for all samples (B) Comparison of DMS-seq data with MNase-seq coverage along the adenovirus genome. Top panel: log fold change of DMS methylation sites of viral samples against free DNA (magenta and blue lines; Fig. 1C). Bottom panel: MNase-seq coverage (according to [43]; blue bars; Fig. 4C)

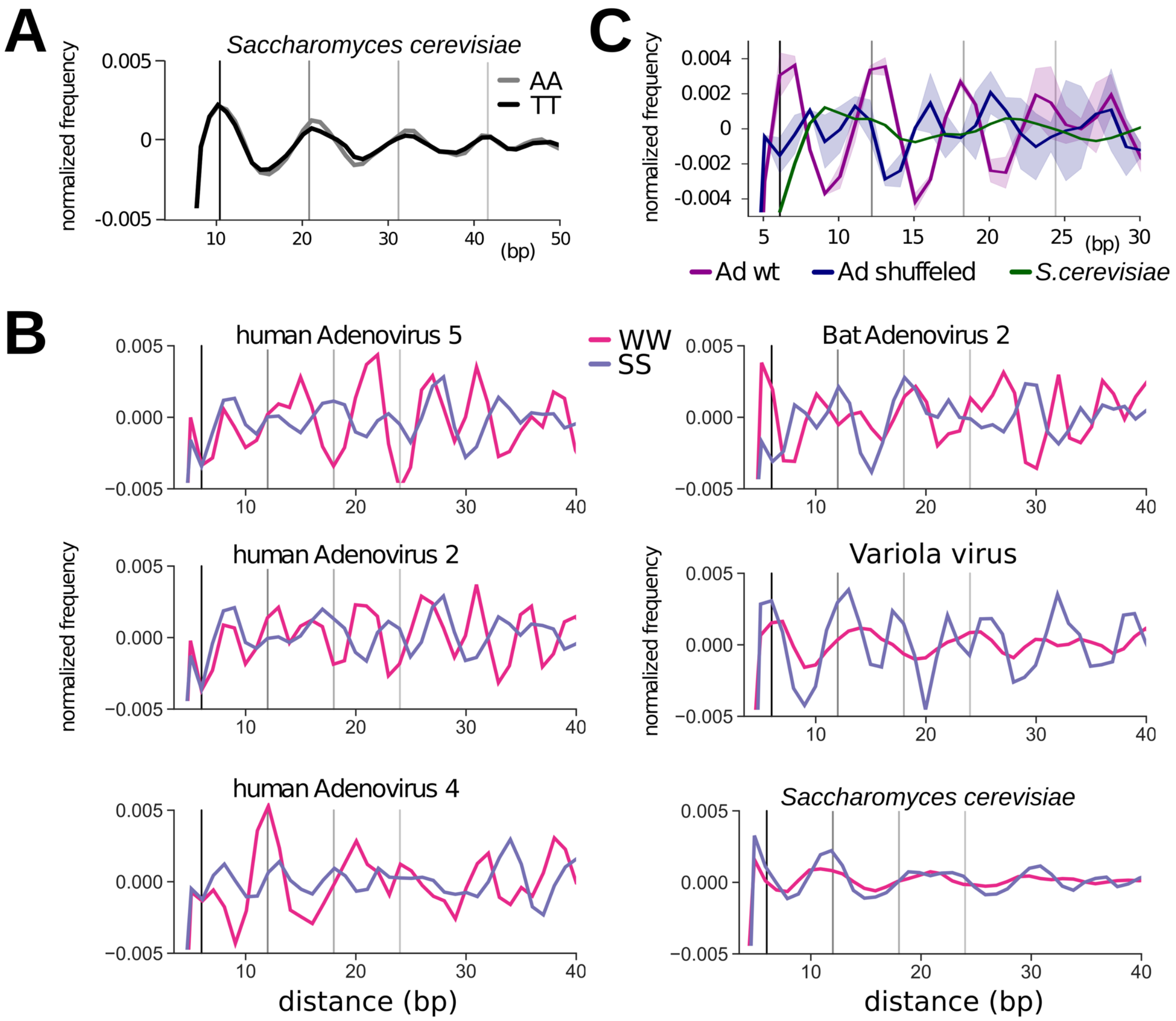


**Figure S4: Dinucleotide autocorrelations**. (A) Nucleotide repeat patterns of ~11-bp AA and TT on the s. cerevisiae genome. Vertical lines are placed every 10.3 bp at the maximum of the autocorrelation curves. (B) SS (G/C) and WW (A/T) di-nucleotide sequence autocorrelations on genomes from different adenovirus species, including Variola virus and the yeast genome. Vertical lines are placed every 6.1 bp. (C) Di-nucleotide (~6-bp) periodicity of the T/C (YY) autocorrelation on the human Adenovirus 5 genome. The same analysis was performed with the shuffled Adenovirus genome and the *s. cerevisiae* genome. Vertical lines are drawn every 6.1 bp.

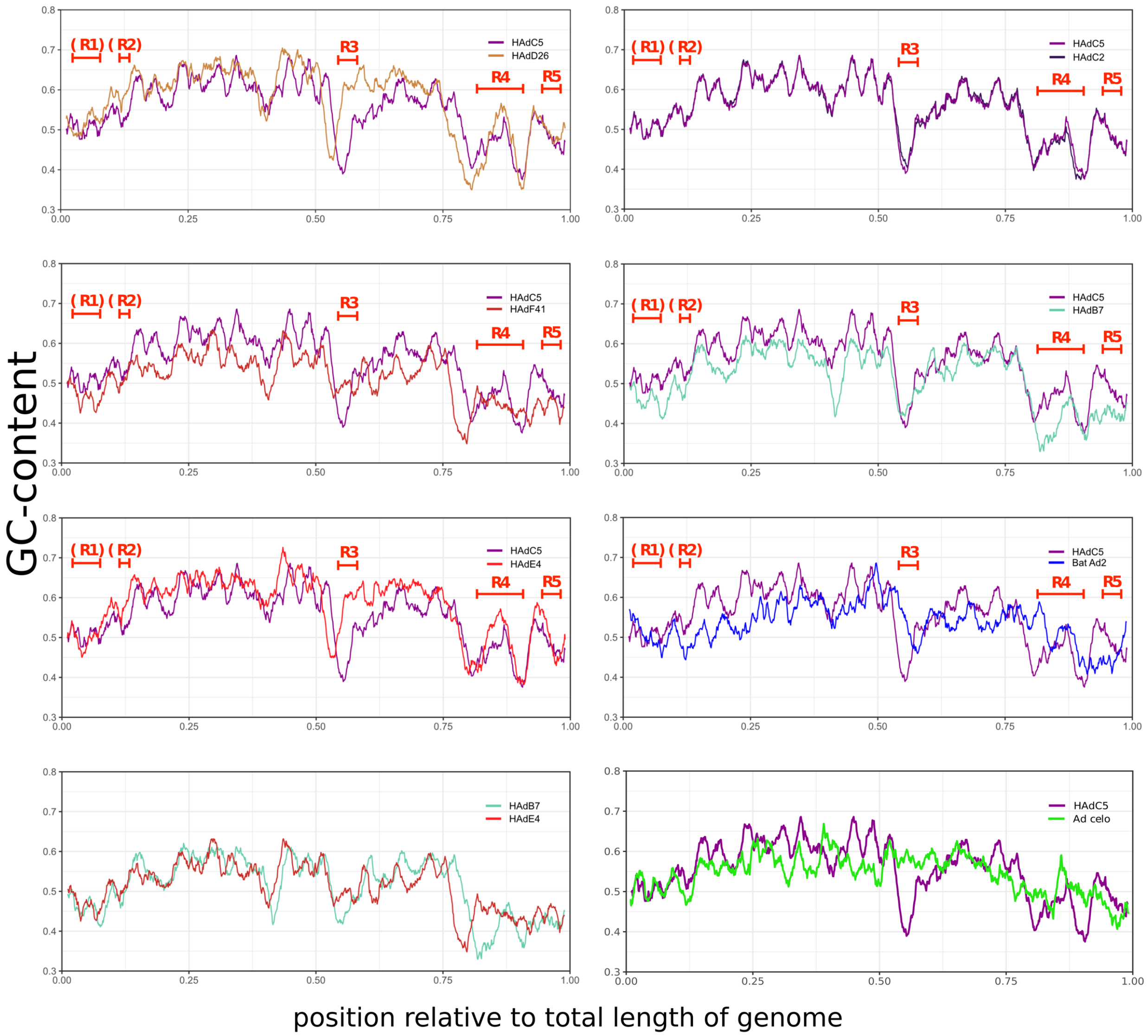


**Figure S5: Comparisons of GC-content distribution along various adenoviral genomes.** Y-axis shows the GC-content. Because of the variable sizes of tested genomes, the X-axis represents the relative positions along the entire length of the genome.
